# Multiplexed Digital Characterisation of Misfolded Protein Oligomers via Solid-State Nanopores

**DOI:** 10.1101/2023.08.09.552642

**Authors:** Sarah E. Sandler, Robert I. Horne, Sara Rocchetti, Robert Novak, Nai-Shu Hsu, Marta Castellana Cruz, Z. Faidon Brotzakis, Rebecca C. Gregory, Sean Chia, Gonçalo J. L. Bernardes, Ulrich F. Keyser, Michele Vendruscolo

## Abstract

Misfolded protein oligomers are of central importance in both the detection and treatment of Alzheimer’s and Parkinson’s diseases. However, accurate high-throughput methods to identify and quantify oligomer populations are currently lacking. We present here a single-molecule approach for the detection of oligomeric species. The approach is based on the use of solid state nanopores and multiplexed DNA barcoding to identify and characterise oligomers from multiple samples. We study α-synuclein oligomers in the presence of several small molecule inhibitors of α-synuclein aggregation, as an illustration of the applicability of this method to assist the development of diagnostic and therapeutic methods for Parkinson’s disease.

## Introduction

The presence of misfolded protein oligomers is associated with the onset and progression of several neurodegenerative disorders, including Alzheimer’s and Parkinson’s diseases^1,2^. These species are likely to form because many proteins may be present in the cell at supersaturated concentrations, making them prone to aggregation, and driving the interconversion between functional states and aberrant self-assembled multimerised states^3,4^. To prevent this outcome, under normal conditions, the protein homeostasis systems including molecular chaperones and the ubiquitin-proteasome and endosomal-lysosomal degradation pathways^5,6^ ensure the correct folding and complexing of proteins and removal of aggregates^7,8^. The metastable proteome becomes increasingly unstable however, as the body ages and experiences stresses, concomitant with these maintenance pathways becoming less efficacious^9,10^. This leads to uncontrolled protein aggregation and the accumulation of these misfolded oligomers, eventually converting to highly ordered polymeric fibrils^1,2,11^.

Numerous neurodegenerative diseases are thought to result in part as a result of this, as aggregates accumulate and interfere with crucial neuronal functions^1,2,12,13^. The aggregation of α-synuclein (αS) for example, is associated with the initial neurodegenerative processes underlying Parkinson’s disease, in which αS aggregates, and misfolded oligomers in particular, exhibit various mechanisms of cellular toxicity^14–19^. Therapeutic efforts directed at this area have not yet resulted in approved drugs^20^, however, in part because they are based on readouts related to αS fibrils, which are the endpoint of the aggregation process. These highly ordered structures are thought to be largely inert in terms of neuronal toxicity, although they can catalyse the formation of further oligomers via a process termed secondary nucleation^21–26^ (**Figure 1A)**. To date, most investigations into the aggregation process rely on detection of fibrils using amyloid binding dyes, such as thioflavin T (ThT), that fluoresce strongly upon binding to fibrils. This approach, however, does not provide a direct measure of the oligomers present, the population of which vary according to the mechanism of aggregation^27,28^.

**Figure 1.**
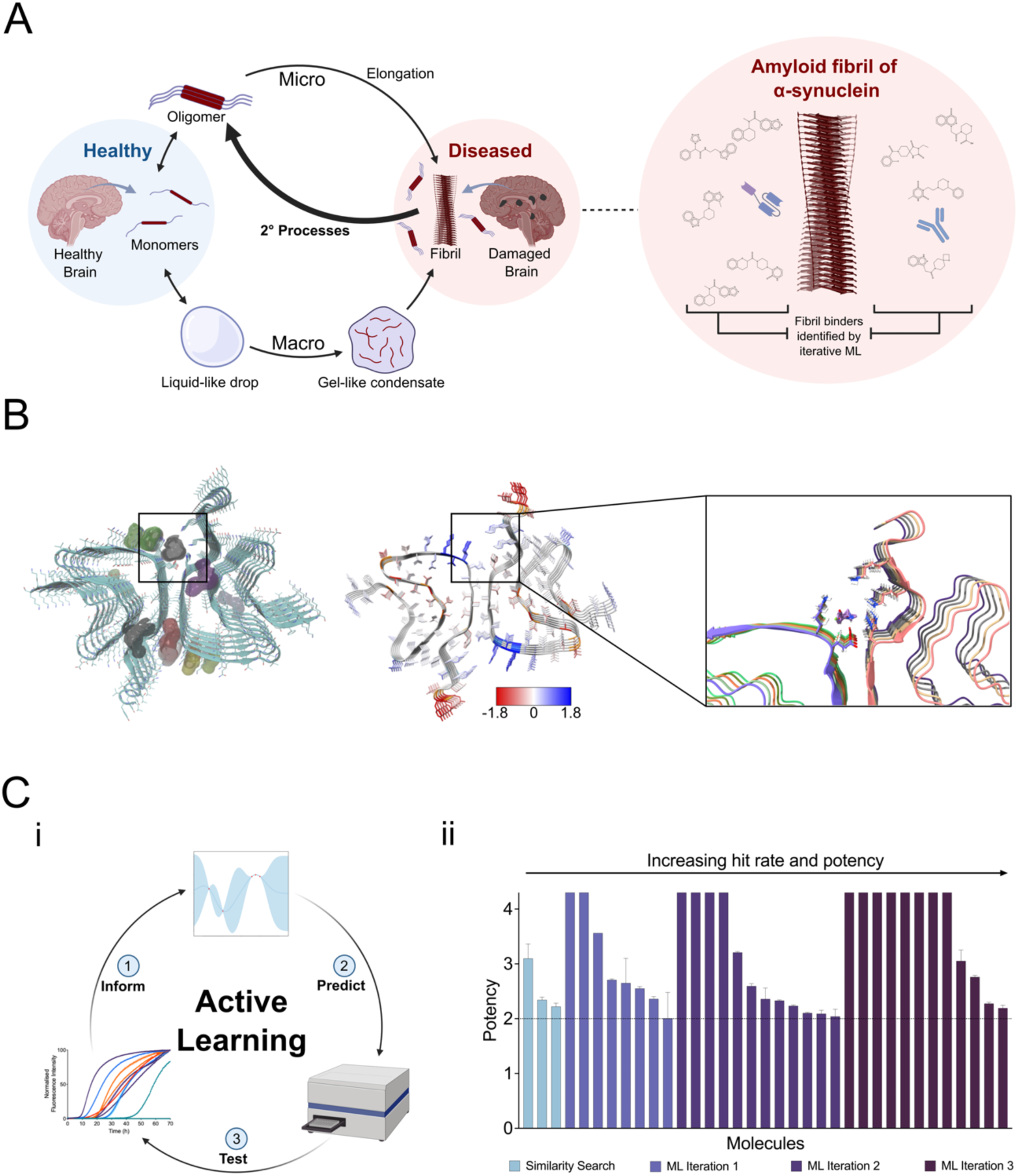
αS aggregation in Parkinson’s disease and its inhibition by small molecules. This drug discovery approach is from parallel work by the authors^38^. (**A**) The age-related progressive impairment of the protein homeostasis system leads to the aberrant misfolding and aggregation of αS into toxic oligomeric species, which eventually convert to amyloid fibrils. These fibrils are observed as the primary constituent of Lewy bodies, a hallmark structure observed in brain cells of patients suffering from the disease. Fibrils can act as a catalyst for further oligomer formation via secondary processes such as secondary nucleation from catalytic sites on the fibril surface and fragmentation of the fibrils into smaller species. Secondary processes are the key generator of oligomeric species. **(B)** A structure-based iterative machine learning strategy comprised of docking simulations followed by cycles of active machine learning was employed to identify secondary nucleation inhibitors. The former consisted of docking small molecules to the catalytic aggregation sites on the fibril surface, identified in the cryo-EM structures of the fibrils via pocket identification software^51^, surface solubility predictions (colour coded from low to high, red to blue)^52^ and experimental knowledge. **(Ci)** Iterative active machine learning was applied to a set of experimentally-validated hits, initially identified via docking, and their analogues. **(Cii)** This successively improved molecule potency with every iterative experimental screen, as the information from each screen was fed back into the active learning algorithm. The 8^th^ hit from iteration 3, I3.08, is used as a tool compound here.

A promising therapeutic strategy is the blocking of secondary nucleation, which is a key accelerator of oligomer production^29,30^. Specifically targetting oligomer producing steps is essential. If fibril elongation were to be inhibited for example, this would slow the formation of endpoint fibril but increase the population of misfolded oligomers by shifting the aggregation pathway more strongly toward secondary nucleation^27^. Previous work has shown methods of isolating specific mechanisms of aggregation and their respective rates experimentally, and subsequently inferring the oligomer populations at a given time via fitting to an analytical model of the aggregation process^21,28^. Theoretical predictions were previously experimentally validated by taking samples during the aggregation process, tracked via ThT, and separating by size exclusion chromatography (SEC) before measuring the monomer equivalent oligomer concentration in each lyophilized sample via mass spectrometry or ELISA^28^. While this is a valid strategy, it is hampered by low throughput and technical challenges in the implementation. Therefore there remains a need to experimentally probe the oligomer population in a non-disruptive and higher throughput manner to determine the size distributions of the oligomer population over time at single particle resolution^31^.

Thus far, single-molecule techniques using fluorescence have shown promising results in characterizing oligomer distributions^31^. For example, confocal two-color coincidence detection (TCCD)^32^, fluorescence correlation spectroscopy (FCS) measurements^33^, single molecule total internal reflection fluorescence (TIRF) imaging^34^, single-molecule spectrally-resolved points accumulation for imaging in nanoscale topography (sPAINT)^35^, atomic force microscopy (AFM)^36^, and micro free flow electrophoresis (µFFE)^37^ have all allowed study of oligomer distributions under near physiological conditions. Additionally, it has been shown that using µFFE one can ascertain oligomer populations in the presence of specific secondary nucleation inhibitors^37,38^. One major limitation of these approaches, however, is the low throughput at which molecules can be tested.

A promising alternative towards achieving high-throughput is nanopore sensing, a single-molecule technique which relies on applying an electric field to drive molecules through a nanosize opening, allowing one to measure changes in ionic currents relating to the size, shape and charge of the molecule entering, or translocating, through the pore^39^. Broadly speaking, there are two types of nanopores, biological, based on using a porous protein, and solid-state, which involve fabricating materials to create a nanosized opening. Platforms containing biological nanopores are commercially available from Oxford Nanopore Technology. However, due to size, these are mostly restricted to DNA sequencing^40^ or rely on protease cleavage of samples before nanopore measurements^41^. Recently, the ability to discriminate between alpha-synuclein variants has been accomplished using biological nanopores^42^. However, using solid-state nanopores eliminates the need for fragmentation and allows the size of the nanopore to be directly tuned and optimized for detection of the analyte of interest. Auspiciously for potential high throughput applications, it has recently been demonstrated that solid-state nanopores could be manufactured at scale^40^.

Previously, solid-state nanopores have proven to be a useful tool for the detection of proteins^43^, as well as a way to study their conformations and interactions^44^. One of the major challenges associated with studying proteins in solid-state nanopores, however, is the speed at which they translocate. However, this challenge can be overcome with approaches such as employing bilayer-coated solid-state nanopores^45^ or by increasing the current bandwidth which increases the time resolution of the measurement^46^. In one case, αS oligomerization was even studied in solid-state nanopores using a Tween-20 coating. While these approaches are effective for studying single proteins, they are not easily adapted for multiplexed sensing.

Since it has been demonstrated that the combination of solid-state nanopores and digitally-encoded DNA nanostructures allows for highly multiplexed detection of single molecules^47,48^, in this work, DNA nanostructures are used to study the effect of small molecule inhibitors of αS secondary nucleation in a multiplexed assay. The small molecule inhibitors were determined via parallel work done by the authors^38,49^. In this work, inhibitors were initially identified via in silico docking to a putative catalytic site that promotes oligomer formation (**Figure 1B**) on the surface of αS fibrils followed by optimisation in aggregation assays via active machine learning (**Figure 1C**)^38,49,50^.

Application of nanopore detection to quantitative protein oligomer analysis therefore offers another useful application of this technique, with the potential of high throughput analysis of a challenging target and an associated benefit to therapeutic programmes targeting these misfolded protein aggregates.

## Results

### DNA nanostructure design for oligomer capture

A DNA nanostructure was designed that could couple to azide-tagged αS aggregates and uniquely identify them (see Methods and **Figure 2**). Using a single-stranded DNA (ssDNA) backbone as a scaffold, complementary staple DNA oligonucleotides were combined with additional oligonucleotides for detection and digitization in a one-pot reaction and annealed. DNA dumbbells allowed digitization of the structure. Their presence created a structured spike in the nanostructure, while their absence left a flat spacer region, corresponding to either a ‘1’ or ‘0’. In the proof of concept presented here, only five spike/spacer regions were used, allowing for 2^5^ (32) combinations of barcodes. This design was based on previous work and was optimized to create clearly distinguishable spikes in nanopores of ∼15 nm diameter^53^. However, this has the potential to be expanded with further optimization, as we have previously experimentally demonstrated that one can fit 56 bits onto a single DNA carrier, allowing for a library of 2^56^ (>10^16^) molecules^54^. Another section of the nanostructure contained two DNA strands, one 21 base pair (bp) sequence labelled with a dibenzocyclooctyne (DBCO) tag and one which had partial complementarity to both the backbone and the sequence containing the DBCO, connecting the DBCO tagged region to the rest of the nanostructure (**Figure 2A**).

**Figure 2.**
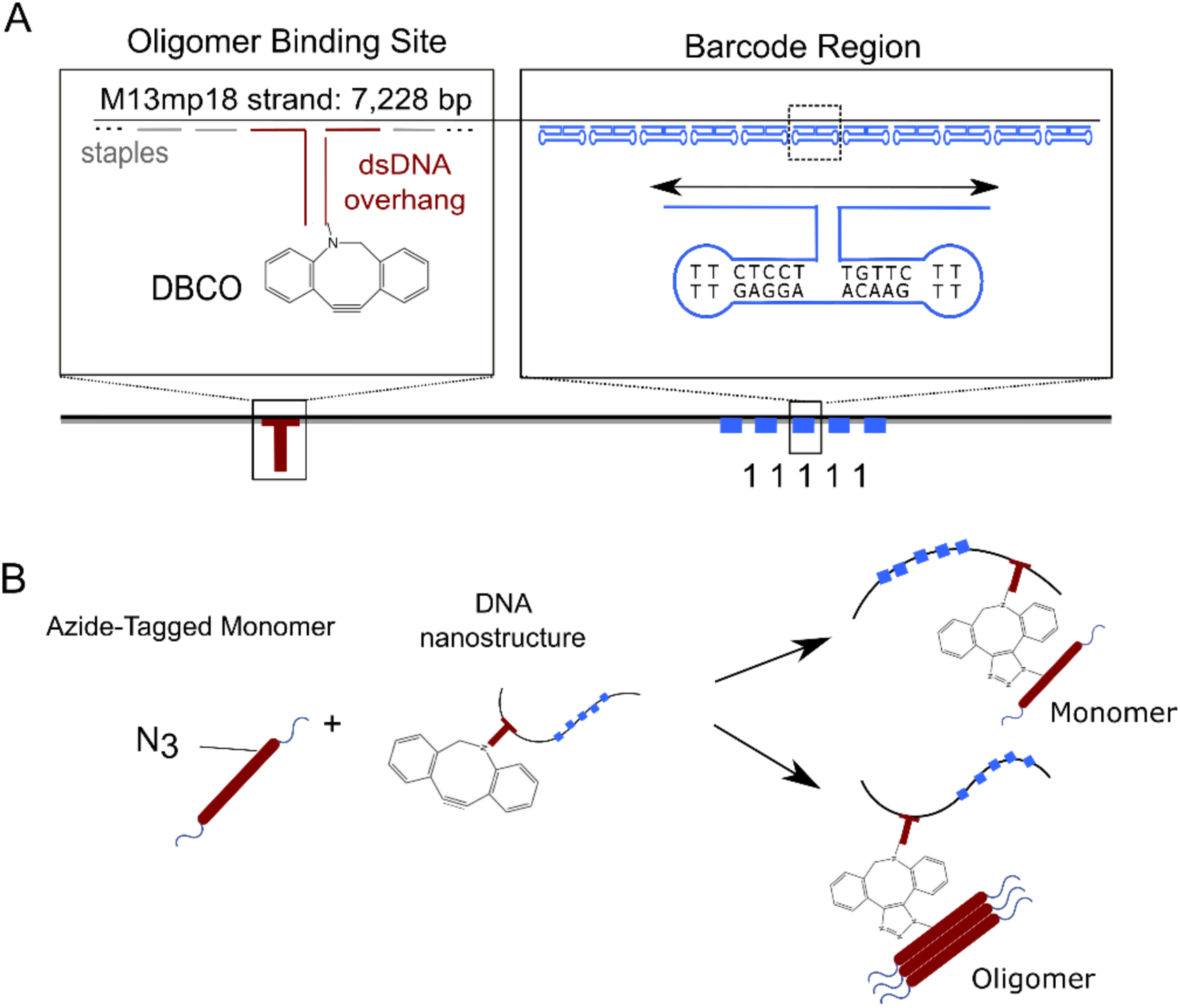
Design of a DBCO-DNA nanostructure for the capture of azide-labelled αS aggregates. (**A**) Schematic of the DNA nanostructure containing the DNA barcode region and a DBCO-tagged dsDNA overhang for click coupling to azide-tagged N122C-αS. DNA barcodes allow for a digital readout of the single-molecule translocations using DNA dumbbells to create distinct 1 or 0 bits. **(B)** N122C-αS is tagged with iodoacetamide-PEG_3_-azide and then incubated with the DBCO-tagged nanostructure, allowing facile click coupling of the two components.

The DBCO-labelled nanostructure was then combined with azide-tagged N122C αS samples for click coupling and subsequent detection (**Figure 2B**). The azide-tagged N122C αS monomer was prepared via reaction of the reduced cysteine thiol with the iodoacetamide moiety of iodoacetamide-PEG_3_-azide. This reaction was monitored until completion via LCMS (**Figure S1**). The monomer was isolated via SEC before use in aggregation experiments and subsequent coupling to the DNA tags.

### Detection of stabilized oligomers via DNA nanostructures and nanopores

We chose to first test the ability of the nanopores to act as a device to detect oligomers using a stabilised oligomeric species. Stabilised αS oligomers have been extensively characterised previously^55,56^. They are typically obtained using methods such as hyper-concentration and lyophilisation, and as such have limited physiological relevance. However, they do offer a useful test case for oligomer detection methods due to their greater stability, higher concentration and larger size^56^. Stabilised oligomers were used to optimise coupling times to the DBCO-tagged DNA barcodes and also to test whether an appreciable difference could be observed in monomeric and oligomeric samples in the nanopore. Successful click coupling of the samples was confirmed via PAGE (**Figure S2**, **Table S1**), where monomer-bound DNA was observable.

Samples of the coupled DNA-protein assemblies were pushed through a nanopore using an electric current as a driving force (**Figure 3A**). The negatively charged nanostructure aided insertion into the pore when a current was applied. In this case, since the protein was also negatively charged at the pH used, the translocation was sped up. As the structures translocated through the nanopores, they created unique signals (**Figure 3B**). Monomer samples were compared against stabilized oligomer samples. Because the molecular weight of monomeric αS is ∼14 kDa, and as can be seen from the low percentage of additional spikes on the nanostructure from the monomer sample in **Figure 3C**, we can assume it is too small to be observed via the 15 nm nanopore. In this experiment, the samples containing no protein, only monomer, or stabilised oligomers were tested in different pores as a control to ensure there were no inter-sample interactions. The lack of events observed in the pure monomer sample allows us to clearly distinguish the samples with and without oligomers by their current traces, and removes the monomers as a source of noise as their signal is too low to be detected in a nanopore of this width. This demonstrates how the customizable dimensions of solid state nanopores can be utilized to focus on the subsample of interest. A significant difference in the percentage of events with proteins attached to the DNA barcodes was observed between the oligomeric and monomeric samples, demonstrating the potential utility of the approach to determining oligomer levels in a sample (**Figure 3C**).

**Figure 3.**
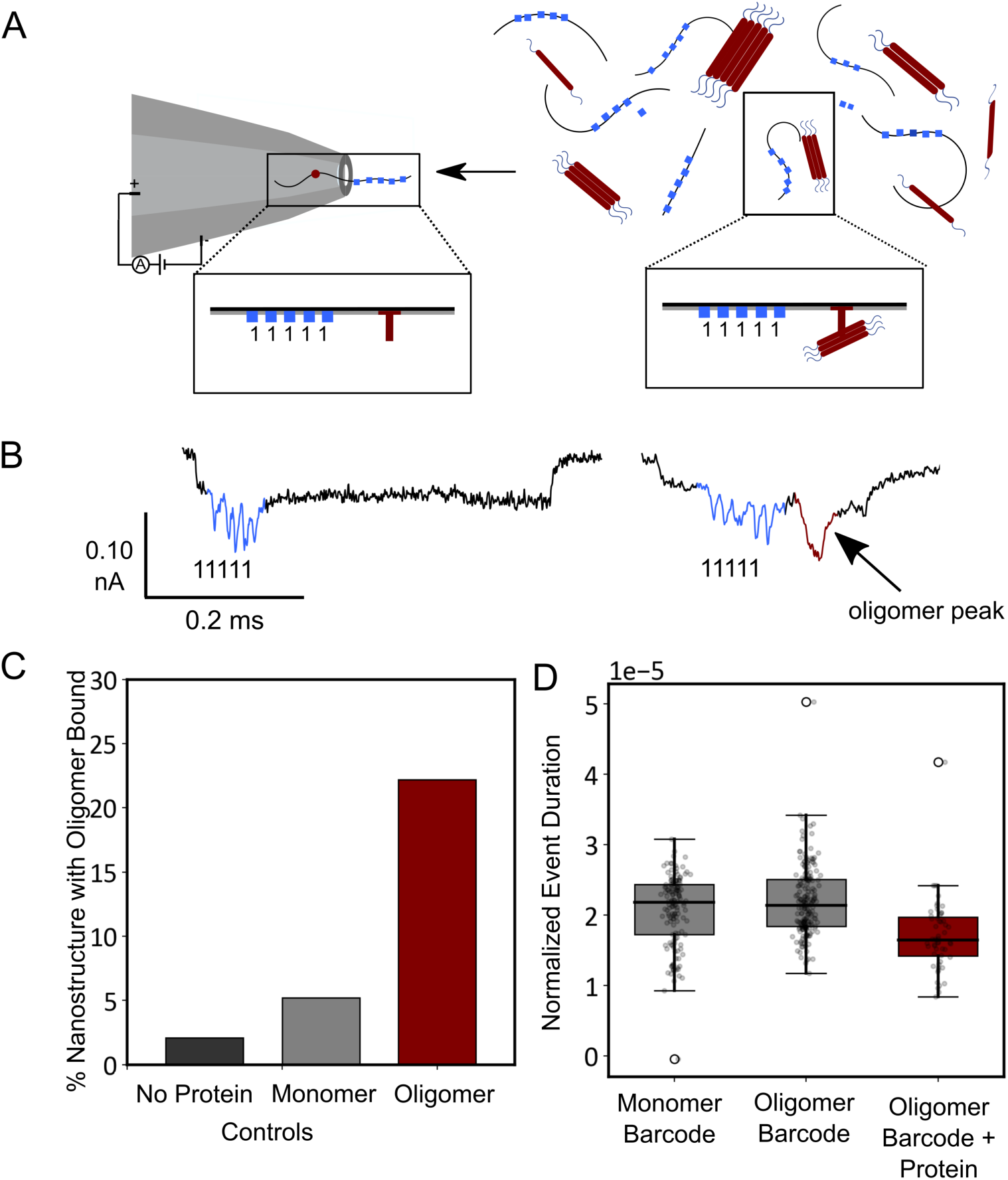
Detection of stabilized oligomers using nanopores. (**A**) Nanopore schematic representing the nanostructures with and without oligomers bound. **(B)** Current trace of nanopore with no protein bound (left) and with an oligomer bound (right). **(C)** Percentage of events with spike for control sample without protein added (N=48), monomer only added (N=154) and stabilized oligomer added (N=248). The samples with just monomer and stabilized oligomer act as controls and show the low percentage of false positives. **(D)** Normalized event duration (normalized to pore baseline current) for samples with barcode only or with barcode and protein spike mixed with either monomer or oligomer.

It should be noted that the oligomeric samples also contained a significant proportion of monomer, which is otherwise challenging to separate entirely from the oligomer sample. Of the observed events, ∼22.2% had a protein oligomer peak attached to the DNA nanostructure. The rest of the events exhibited no peak due to being bound to monomeric protein, which makes up the majority of the sample. The ability to measure with this background present is essential given the additional time cost and potential bias introduced by the need to separate oligomeric species from the bulk monomer. These events can be separated both by observing the nanopore signal generated, where little to no protein peak signifies either an uncoupled DNA nanostructure, or a nanostructure coupled to only monomer, as well as by using parameters such as event duration (**Figure 3D**). Because the protein is negatively charged, the event duration decreases in samples with bound proteins. As these samples were measured in different pores at different times, to ensure no cross sample contamination and reliable controls, the duration also must be normalized to the baseline current (I_0_), thus a normalization was carried out as explained in the methods (**Equation 2**).

### Effect of inhibitor molecules on the oligomer production

Having optimised the conditions, we then tested more challenging “on time-course” samples. We carried out an aggregation beginning from monomer, under conditions designed to promote secondary nucleation^21,57^. This assay has been fully characterised for an AlexaFluor-488 tagged N122C vs WT in previous works, and azide tagging did not substantially alter this behaviour^37,57,58^. Oligomer populations in this scenario are significantly lower in concentration compared to the stabilised oligomer case, and they are transient. On time-course samples of αS are only stable for ∼24 h post extraction, compared to αS stabilised oligomers which persist for up to a week after production if left at room temperature.

The on time-course experiment was designed to better mimic the processes and species that may occur in vivo. In order to induce αS aggregation via secondary nucleation in vitro at neutral pH, a small amount of pre-formed seed was added (100 nM monomer equivalents) in the presence or absence of aggregation inhibitors of interest (**Figure 4A,B**). The aggregation process was followed using ThT flourescence. The 3 samples of interest were a control containing only 1% DMSO, another control containing Anle-138b^24^ (an αS aggregation inhibitor that entered clinical trials but exhibited only mild efficacy) in 1% DMSO, and a small molecule identified previously via structure-based machine learning methods, I3.08, also in 1% DMSO. DMSO was used to dissolve the molecules before adding to the aqueous protein sample.

**Figure 4.**
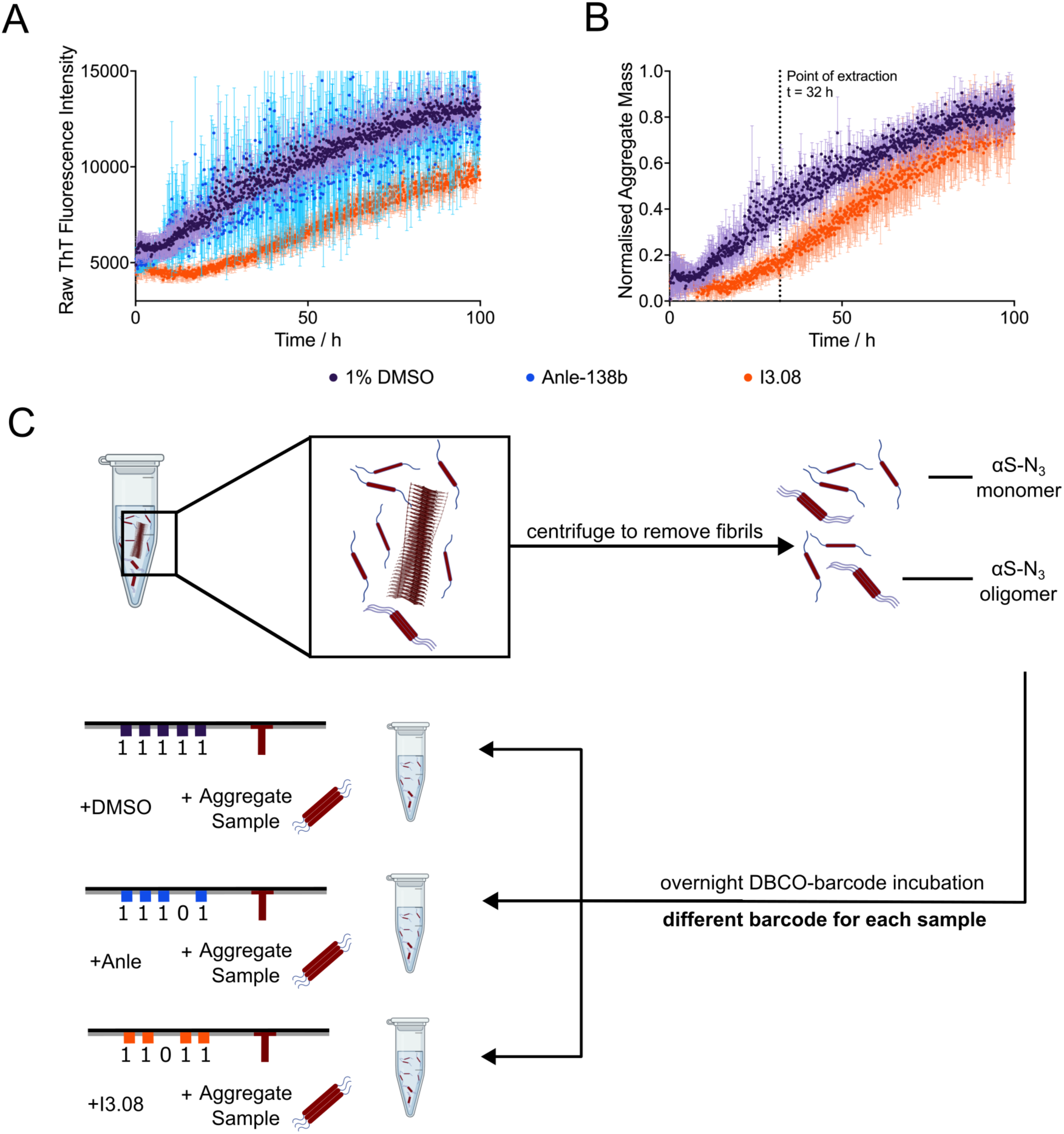
Preparation of on time-course oligomeric samples in the absence and presence of inhibitor molecules. (**A,B**) Kinetic traces are shown of a 10 µM solution of azide-tagged N122C-αS supplemented with 100 nM pre-formed seeds (pH 7.4, 37 °C, shaking at 200 rpm, error bars denote SD) in the presence of 1% DMSO (purple), 25 µM Anle-138b (blue) or I3.08 (orange). The raw fluorescence (A) and normalised fluorescence (B) are shown. The endpoints were normalised to the αS monomer concentration at the end of the experiment, which was detected via the Pierce™ BCA Protein Assay at *t* = 100 h. Anle-138b could not be suitably normalised due to the noise of the sample. **(C)** Samples were extracted at 32 h from the time course of aggregation and centrifuged to remove fibrils from the mixture, leaving only αS monomers and soluble oligomeric species for analysis. These samples were then incubated with a unique DBCO-tagged DNA barcode overnight before analysis via solid state nanopore detection.

Our previous work reports that I3.08 binds to the fibrils, not the monomer or oligomers, and in so doing blocks autocatalytic aggregate formation^38^. Fibrils are removed prior to nanopore measurement by centrifugation so only the oligomer and monomer population remain. The molecular mechanism of Anle-138b has not been published, and is presumably not known as it has mainly been characterised in mouse models. Our work indicates it may operate by a similar mechanism only in a less effective manner. The aggregation was accelerated via shaking, which was necessary to complete the aggregation under cellular buffer conditions in an experimentally accessible time frame, but created a more challenging paradigm for the inhibitors to function in. The inhibitors are only capable of preventing aggregation via secondary nucleation, not via fragmentation resulting from mechanical shearing. Nonetheless a significant inhibition of fibril accumulation was still observed for inhibitor I3.08, although not for the control inhibitor Anle-138b. Samples were then taken mid-way through the time course to determine whether a similar reduction in oligomeric species was also observed.

Samples were extracted at 32 h into the aggregation time course and centrifuged to remove fibrils before click reaction of the azide tagged αS with unique DBCO-tagged DNA barcodes overnight at a ratio of 1:1 (DBCO: initial monomer concentration) (**Figure 4C**). The aggregation reaction was diluted 2500-fold for this coupling, effectively quenching further aggregation. In the absence of conditions favouring phase separation^59^, αS does not continue to aggregate under experimentally accessible timescales at concentrations below 5 µM regardless of the conditions^25,60,61^. Each sample was labelled with a different DNA barcode; DMSO (11111), Anle-138b (11101) and I3.08 (11011). Additionally, DBCO/N122C-azide required at least > 3 h incubation time for the reaction to proceed significantly (**Figure S3**). The rate was tested by sampling 1, 3 and 12 h incubation times. No observable shift in PAGE was visible for 1 or 3 h, but an observable shift was visible for the sample incubated for 12 h. These results demonstrate that we can multiplex the samples without concern for significant further coupling reactions from any residual unreacted azide/DBCO species during the nanopore measurement. Concerns over possible interchange of monomers in the sample between oligomers of different samples were addressed by the dilution at this stage, with the expectation that interactions become essentially unfeasible. Additional repeats were done by mixing duplexed DMSO and I3.08 samples (**Figure S4**). Similar results for samples tested in duplex and triplex support this assumption.

### Multiplexed digital nanopore read-out of the effect of inhibitor molecules

Using the method described above, the samples were then run through the nanopore. Analysis of both the number of events containing a discernible DNA barcode and an attached protein peak, and the area of the protein peak, showed a change in oligomer distribution compared to the DMSO control (**Figure 5**). The DNA barcode is the observable quantity, and so a ratio of the barcode with bound oligomer vs unbound was calculated as described in the Methods. The nanopores were fabricated to be 10-15 nm, such that monomeric proteins would not be observable, while oligomeric species would be observable. The DMSO barcode was 29.8% bound to protein oligomers, the Anle-138b sample was 41.8% bound and the I3.08 sample was 14.4% bound (**Figure 5B**). The size distribution of the oligomers broadly matched this trend, showing decreasing oligomer mass from the DMSO sample to the Anle-138b sample, which contained a large number of small oligomers as explained below, and lastly the I3.08 sample (**Figure 5C**). This was calculated using **Equation 3**. As the samples were run simultaneously in the same pore, no normalization was required. These results show that compound I3.08 reduced oligomer production relative to the untreated control, and that it was a better inhibitor of oligomer production than Anle-138b.

**Figure 5.**
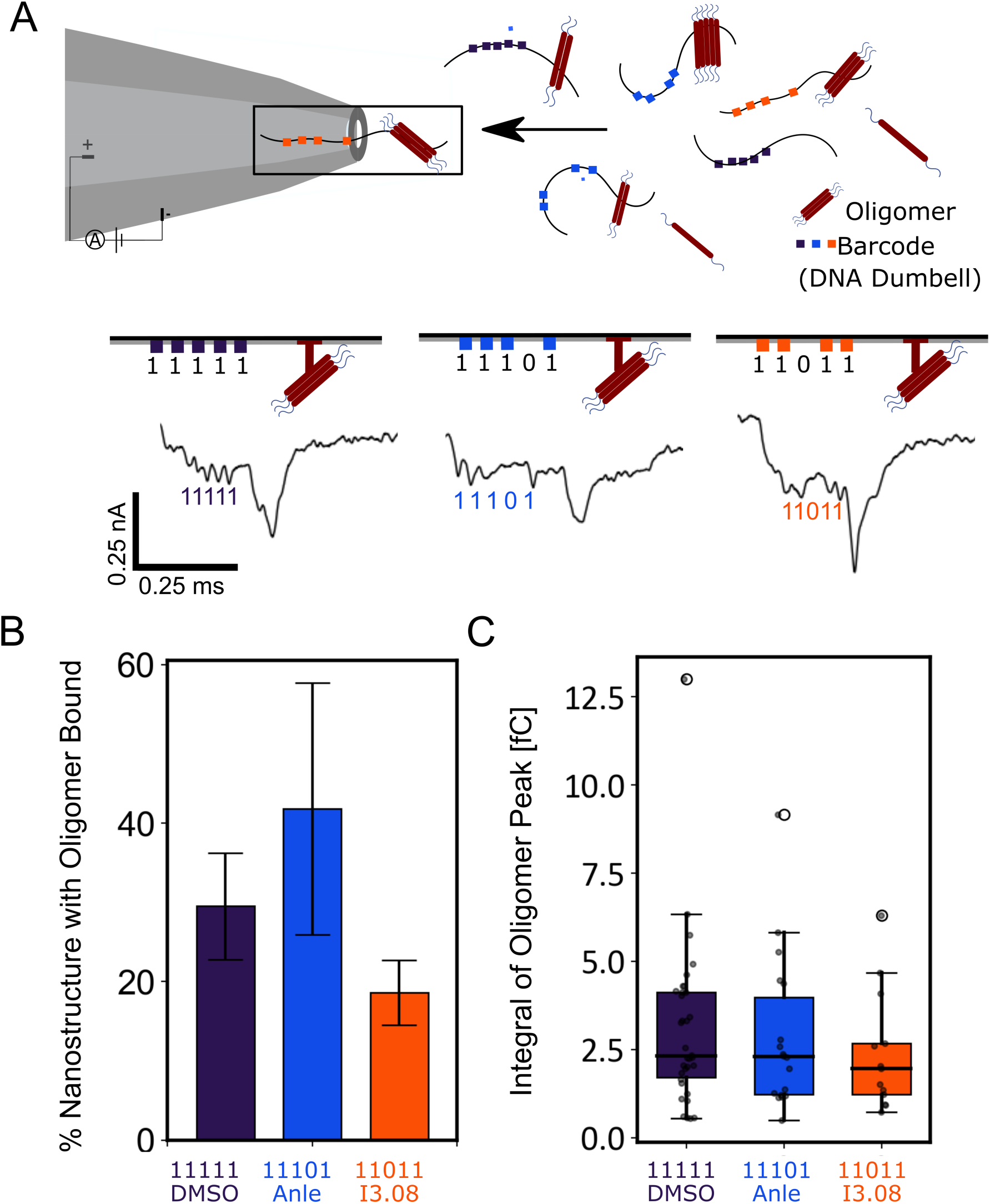
Schematic of the multiplexing pipeline and comparison of two different inhibitor molecules effects against on time-course samples. (**A**) Samples are tagged with a unique DNA barcode that allows identification in a multiplexed mixture, increasing the throughput. The events observed as the oligomers translocate through the nanopore can then be analysed to give an oligomer number per tag, and a relative area under the curve of each tag, proportional to oligomer size. (**B**) The fraction of events with an oligomer bound to the DNA barcode; DMSO (purple) (N=114±7), Anle-138b (blue) (N=43±16) and I3.08 (orange) (N=90±4). The standard deviation comes from repeats where the samples were combined, diluted in measurement buffer and measured for ∼1 h. (**C**) Area of the current drop of the protein spike caused by bound oligomer in the DMSO (purple), anle-138b (blue) and I3.08 (orange) samples. A larger area implies larger species are bound to the barcode on average.

Interestingly, first I3.08 and DMSO were tested in duplex and a similar baseline level noise (∼6 pA) was maintained throughout the measurement. With the addition of Anle-138b in triplex with the other samples, the noise level increased (**Figure S5**). This is consistent with the kinetic data (**Figure 4A**). The Anle-138b sample exhibits a noisy kinetic trace, consistent with increased formation of particulates, and has a correspondingly greater oligomer population. The increase in nanopore noise is most likely due to the larger oligomers present rapidly translocating through the pore at the beginning of the measurement. After 3 min, most of the larger oligomers have translocated through the pore which leads to the baseline current and noise resuming back to normal. This is also consistent with the number of events measured for Anle-138b (N=43), where fewer discernible events with Anle-138b barcode 11101, as compared to DMSO barcode 11111 (N=114) and I3.08 barcode 11011 (N=90) were observed despite all samples being added at equal concentration.

### Comparison with a micro free flow electrophoresis (µFFE) method

For comparison, a state of the art technique in protein oligomer detection is micro free flow electrophoresis (µFFE), which allows full characterisation of the oligomer distribution in physiological conditions, and has been previously applied to ascertaining oligomer populations in the presence of a closely structurally related inhibitor to the one used here^37,38^. While µFFE requires insoluble fibrils to be removed via centrifugation, no further separation is required, as the technique separates the monomeric fraction from the oligomeric fraction in situ using an electric field across the particle stream that deflects particles based on their electrophoretic mobility. The only disadvantage is relatively low throughput. In that work, a molecule (I3.02) induced a 37% delay in half time of aggregation compared to the negative DMSO control. As a result, there was a 75% reduction in the mass of oligomers present at the half time of the negative control. The aggregation kinetics were carried out under similar conditions as used here, the primary difference in that work being the higher concentration of αS monomer and the molecule (100 µM αS, 50 µM molecule). In this work, molecule I3.08 induced a 57% delay in relative half time and, as measured by nanopore detection, the drop in oligomer events observed was 48% and the drop in oligomer mass was 22% compared to the negative DMSO control. Anle-138b was shown to have lower effectiveness in terms of oligomer number reduction and oligomer mass reduction via both techniques, and so the ranking of effectiveness between nanopore detection and µFFE is in agreement.

The strategy here was to create a novel screening approach for aggregation inhibitors, not to fully characterise the aggregation time course, though this would represent a valid application of the technology. This has however been done multiple times previously^21,57,58^ while oligomer inhibitory screening assays are scarcer, due to difficulty in applying existing methodologies with low throughput. The comparison between µFFE, one of the methods used to carry out a full time course characterisation^37,58^ and then to characterise inhibitor potency^38^, and the nanopore method shown here, demonstrates that both are effective at ranking molecules in terms of molecule potency.

## Discussion

We have reported a nanopore detection method for misfolded protein oligomer detection and analysis, with a detection limit on par with current state of the art techniques, but with significantly greater potential for throughput. To illustrate the method, we applied it to detect the inhibition of αS oligomer production by small molecules in clinical development. This result was obtained with the additional benefit of multiplex capability and higher throughput.

While the nanopore system has many advantages, there are also some drawbacks. A drawback of the large dilution step required for measurement in the nanopore is the possibility that some of the oligomers may dissociate during the DBCO coupling step (12 h) due to the large dilution (2500 fold). This is a feature of most single-molecule techniques which require low concentrations in order to have clear signal to noise. However, αS is a useful test case in this scenario given that its kinetics are relatively slow and its oligomers are stable^37,62^ over the time scales investigated so we consider the measured sample to be a reasonable reflection of the population present at the extraction stage. In further developments, a cross-linking step could be introduced to ensure that the protein sample extracted exactly matches the one measured. This carries the risk of cross linking separate oligomers (potentially mitigated by appropriate dilution) and adds further processing steps, issues which we sought to avoid in the interests of throughput and preventing biasing of the oligomer population. Alternatively, if the dissociation rate in a particular case was a cause for concern, a more reactive click pair could be employed than the one used here, or the coupling could be carried out at higher concentration (followed by dilution immediately prior to measurement) to obtain coupling over a shorter time scale and slow dissociation. A restraint on the click coupling reaction is that the sample conditions cannot be altered in terms of pH or temperature as this would affect the oligomer distribution.

An additional concern with the nanopore measurement is the high salt concentration required for measurement, which may perturb the aggregate distribution. However, the click chemistry reaction was performed in PBS, and the samples were only mixed in the detection buffer directly before measurement. The ratio of protein bound to unbound DNA nanostructures also did not change over the time of the observation (**Figure S6**) suggesting this is not a major issue. Again, cross-linking could remove this problem if necessary. In the interests of throughput, however, and for cases where there is a clinical trial benchmark all that would be required is a relative measurement to compare the effect of different inhibitors. As the samples are measured under the same conditions, a ranking of effectiveness can still be obtained. For protein systems that aggregate very rapidly, the concern is more that the monomers and oligomers may further aggregate during the click reaction rather than dissociate. We anticipate that for almost all proteins the significant dilution should quench aggregation to a rate that is negligible over the time span of the coupling reaction.

Finally, using nanopores as a tool to measure oligomers does have a fundamental size limit, in that particles larger than the diameter of the pore and smaller than the resolution limit will not be detected. However, with a degree of prior knowledge, the nanopore diameter can be appropriately tailored to the size distribution of interest, allowing sampling of a representative portion of the population.

The results that we have presented illustrate an approach for investigating protein assemblies that are both transient and at very low concentration. We have applied this method to the scenario of early drug discovery for Parkinson’s disease and synucleinopathies in general, where misfolded αS oligomers are considered to be key to pathology. We also show comparable performance to existing single-molecule techniques, but with greater potential for throughput due to the ability to multiplex and upscale. With the introduction of artificial amino acids bearing azides into in vivo models of disease^63^, this also represents a potential approach for directly quantifying oligomer populations in such models, utilising the biorthogonality of the click reaction employed here. We anticipate that this approach could be of significant benefit to researchers working in the field of protein misfolding diseases and protein multimerization, and in early-stage drug discovery research in general.

## Materials and Methods

### Compounds and chemicals

Compounds were purchased from MolPort (Riga, Latvia) or Mcule (Budapest, Hungary) and prepared in DMSO to a stock of 5 mM. All chemicals used were purchased at the highest purity available.

### Recombinant αS expression

Recombinant αS was purified based on previously described methods^60,61,64^. The plasmid pT7-7 encoding human αS was transformed into BL21 (DE3) competent cells. Following transformation, the competent cells were grown in 6L 2xYT media in the presence of ampicillin (100 μg/mL). Cells were induced with IPTG, grown overnight at 28 °C and then harvested by centrifugation in a Beckman Avanti JXN-26 centrifuge with a JLA-8.1000 rotor at 5000 rpm (Beckman Coulter, Fullerton, CA). The cell pellet was resuspended in 10 mM Tris, pH 8.0, 1 mM EDTA, 1 mM PMSF and lysed by sonication. The cell suspension was boiled for 20 min at 85 °C and centrifuged at 18,000 rpm with a JA-25.5 rotor (Beckman Coulter). Streptomycin sulfate was added to the supernatant to a final concentration of 10 mg/mL and the mixture was stirred for 15 min at 4 °C. After centrifugation at 18,000 rpm, the supernatant was taken with an addition of 0.36 g/mL ammonium sulfate. The solution was stirred for 30 min at 4 °C and centrifuged again at 18,000 rpm. The pellet was resuspended in 25 mM Tris, pH 7.7, and the suspension was dialysed overnight in the same buffer. Ion-exchange chromatography was then performed using a Q Sepharose HP column of buffer A (25 mM Tris, pH 7.7) and buffer B (25 mM Tris, pH 7.7, 1.5 M NaCl). The fractions containing αS were loaded onto a HiLoad 26/600 Superdex 75 pg Size Exclusion Chromatography column, and the protein (≈ 60 ml @ 200 µM) was eluted into the required buffer. The protein concentration was determined spectrophotometrically using ε280 = 5600 M^−1^ cm^−1^. The cysteine-containing variant (N122C) of αS was purified by the same protocol, with the addition of 3 mM DTT to all buffers.

### Azide labelling of αS

αS N122C protein was azide-labelled to enable click coupling to DNA tags. N122C (200 µM, PBS, pH 7.4) was incubated with TCEP-HCl (5 eq) for 1 h at RT. The reduced N122C was then desalted with a 5 mL HiTrap desalting column, (Cytiva, 29-0486-84), and eluted in PBS, pH 7.4, 10 mM EDTA and kept on ice. The extend of the reduction was then established via Ellman’s method, and a sample was taken for LCMS analysis. The protein was then incubated with iodoacetamide-PEG3-azide (10 eq) for 3 h at RT, and samples were taken subjected to QTOF MS/MS analysis with a VION mass spectrometer to ascertain the progress of the reaction (**Figure S1**). Deconvolution was conducted in UNIFI software. Upon reaction completion the reaction mixture was separated on a Superdex 75 10/300 GL column (GE Healthcare) at a flow rate of 0.5 mL/min and eluted in PBS buffer to isolate the monomeric fraction and buffer exchange into PBS. The protein concentration was determined spectrophotometrically using ε280 = 5600 M^−1^ cm^−1^.

### αS seed fibril preparation

αS fibril seeds were produced as described previously^61,64^. Samples of αS (700 µM) were incubated in 20 mM phosphate buffer (pH 6.5) for 72 h at 40 °C and stirred at 1,500 rpm with a Teflon bar on an RCT Basic Heat Plate (IKA, Staufen, Germany). Fibrils were then diluted to 200 µM, aliquoted and flash frozen in liquid N_2_, and finally stored at –80 °C. For the use of kinetic experiments, the 200 µM fibril stock was thawed, and sonicated for 15 s using a tip sonicator (Bandelin, Sonopuls HD 2070, Berlin, Germany), using 10% maximum power and a 50% cycle.

### αS stabilised oligomer preparation and subsequent click coupling

αS stablisied oligomers were produced as described previously^56^. Monomeric αS was dialysed into distilled water overnight at 4 °C, using 3.5 kDa MWCO dialysis membranes. 6 mg of the dialyzed protein was aliquoted into 15 mL tubes, flash frozen in liquid nitrogen, and lyophilized for ca. 48 h at room temperature. To prepare the oligomeric samples, the 6 mg of protein was resuspendes in a total of 500 μL PBS to obtain a final protein concentration of ca. 800 μM. The solution was centrifuged if necessary (1 min, 1000 g) to get rid of bubbles formed during the resuspension process. The protein solution was filtered through a 0.22 μm syringe filter and incubated in 1.5 mL tubes at 37 °C for 20–24 h under quiescent conditions. The resultant protein solution was ultracentrifuged (1 h, 288,000 g) to remove any fibrillar species that may have formed during the incubation period, and the supernatant was removed and retained. Each aliquot of supernatant was passed through four 0.5 mL 100 kDa centrifugation filters sequentially (2 min, 9300 g), in order to remove excess monomeric protein as well as the low levels of very small oligomers. To estimate the total mass concentration of the final oligomeric solution (i.e., total concentration in monomer equivalents), the absorbance was measured at 275 nm, using a molar extinction coefficient of 5600 M^-1^ cm^-1^. This preparation results in an overall oligomeric yield of ca. 1%. Samples were then diluted to a final concentration of 88 nM monomer equivalents in PBS and incubated overnight with a final concentration 4 nM of DBCO tagged DNA nanostructure. The reason this excess was used was to attempt to ensure 1 DBCO tag per oligomer and prevent over tagging (each stabilised oligomer has a reported average monomer count of 22^56^). Subsequent on time course experiments were carried out with 1:1 labelling of the DBCO:monomer given the large excess of monomer:oligomer expected in these samples.

### DBCO DNA nanostructures

DNA constructs with different barcoded regions plus a DBCO labelled overhang sequence were created. Each DNA construct was synthesized from pairing a linearised 7.2 kbp single-stranded (ss) M13mp18 DNA with 40 nucleotide staples complementary to the backbone in order to create a full linearized dsDNA. The backbone and staples are annealed for 45 minutes in a thermocycler. Using a 100 kDa Amicon filter, the sample is then filtered and stored in 10 mM Tris 0.5 mM MgCl_2_ pH 8. The concentration is then measured in a nanodrop spectrophotometer with typical yield ranging from 75-95%. The barcoded region design follows a previous work with dumbbells optimized for read out in 15 nm nanopores^65^. Each ‘1’ bit is made of eleven simple dumbbell hairpin motifs to create the structural spikes which act as a barcode on the DNA nanostructure. This can be optimized to have fewer dumbells per spike if needed. The exact sequences with their numbers are shown in **Table S2** in the Supplementary Information following a previous work^65^. The overhang was created by replacing oligo No. 142 with 51 bp segment containing 30 bp to match the M13 backbone and a 21 bp oligo complimentary to another DNA sequence containing a DBCO label. The 21 bp dsDNA overhang is not large enough to generate a current blockade (an observable signal in the nanopore) which has been confirmed by observation. These sequences can be found in the Supplementary Information **Table S3**.

### Aggregation kinetics and subsequent click coupling

Azide-labelled αS N122C (10 μM) was supplemented with seed (100 nM) under shaking (200 rpm) at 37 °C, PBS pH 7.4 and either 1% DMSO or 25 μM molecule in 1% DMSO. Samples were extracted at the *t_1/2_* of the DMSO sample (30 hours). Fibrils were removed by centrifugation (21,130 rcf, 10 min, 25 °C). Samples were then diluted to 4 nM monomer equivalents in PBS and incubated overnight with 1 eq (relative to initial monomer concentration) of DBCO tagged DNA nanostructure.

### Nanopore fabrication and measurement

The nanopores are made of commercially available quartz capillaries (0.2 mm ID/0.5 mm OD Sutter Instruments, CA, USA). A laser-assisted pipette puller (P-2000, Sutter Instrument, CA, USA) is used to create nanopores with diameters of 10-15 nm. 16 conical nanopores are then placed in a custom templated PDMS chip containing a communal cis reservoir and individual trans reservoirs. In order to generate the current, silver/silver-chloride (Ag/AgCl) electrodes are connected to the cis and trans reservoirs in the PDMS chip. In the baseline buffer solution for the stabilised oligomers (4 M LiCl, 1X TE, pH 8.0) and the on pathway samples (2 M LiCl, 1X TE, pH 8.0), a current-voltage curve is taken in order to estimate the nanopore size. Only one nanopore is measured at a time due to the electronics, thus the trans reservoir contains the electrode with a 500 mV bias voltage and the the central cis reservoir which contains the sample is grounded. The measurement is then run for 1-2 hours until 1500-3000 events are gathered. Typically, of these events, 30% are unfolded and are then analysed.

Current signals are collected using an Axopatch 200B patch-clamp amplifier (Molecular Devices, CA, USA). The set-up is operated in whole-cell mode with the internal filter set to 100 kHz. An 8-pole analog low-pass Bessel filter (900CT, Frequency Devices, IL, USA) with a cutoff frequency of 50 kHz is used to reduce noise. The applied voltage is controlled through an I/O analog-to-digital converter (DAQ-cards, PCIe-6251, National Instruments, TX, US). A LabView program records the current signal at a bandwidth of 1 MHz.

### Nanopore data analysis

The experimental data files are stored as technical data management streaming (TDMS) files from the Labview program recording the raw traces. First, a translocation finder python script is used which identifies the events from the raw traces using user-defined thresholds (minimum 0.3 ms duration, minimum 0.1 nA current drop) and stores them in an hdf5 file. This can be found at https://bitbucket.org/nikaer/nanopyre/. Next, the hdf5 file is loaded into the GUI categorizer python script, found here: https://gitlab.com/keyserlab/nanopycker. Using this, the events are printed and the user can manually sort the events time efficiently into different categories and later print events from the hdf5 file that are assigned to a specific category. In this case the categories were barcode without protein and barcode with protein. The percentage of events with oligomer bound is then calculated using

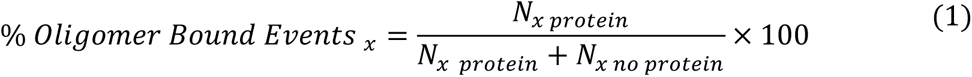

Where *x* is the barcode. This is used in **Figure 3C** and **Figure 5B**. The duration of the events in **Figure 3D** is calculated using

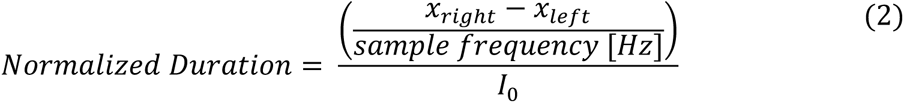

Where *x_right_* is the position of the end of the event and *x_left_* is the position of the end of the event. The sampling frequency is 1,000,000 Hz. *I*_0_is the baseline current because different pores were used for the different measurements with different baselines.

The GUI categorizer is used again on the events with protein to calculate the ECD of the protein peak using

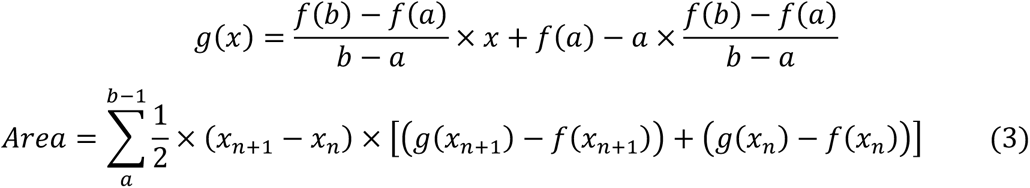

Where *f*(*x*) is the current at point *x*, *a* and *b* are the left and right bounds of the region of interest and *g*(*x*) is the equation of the line connecting *a* and *b*.

### Mass spectrometry

10 µM of preformed αS was incubated with 25 µM of molecule in 20 mM sodium phosphate buffer (pH 4.8) supplemented with 1 mM EDTA overnight under quiescent conditions at room temperature. The supernatant was removed for analysis using a Waters Xevo G2-S QTOF spectrometer (Waters Corporation, MA, USA).

## Acknowledgements

This work was supported by the UKRI (10059436, 10061100). The authors would like to thank ARCHER, MARCOPOLO and CIRCE high performance computing resources for the computer time. S.E.S. acknowledges funding from Oxford Nanopore Technologies, Engineering and Physical Sciences Research Council (EPSRC) and Cambridge Trust. U.F.K acknowledges funding through a ERC-2019-POC PoreDetect 899538. Z. F. B. would like to acknowledge the Federation of European Biochemical Societies (FEBS) for financial support (LTF). The authors are furthermore grateful for financial support from the Cambridge Centre for Misfolding Diseases. The work is supported by the European Research Council (ERC) under Horizon 2020 research and innovation programme PICOFORCE (Grant Agreement No. 883703), THOR (Grant Agreement No. 829067) and POSEIDON (Grant Agreement No. 861950). JJB acknowledges funding from the EPSRC (Cambridge NanoDTC EP/L015978/1, EP/L027151/1, EP/S022953/1). Parts of the figures were created with BioRender.com.

## Supplementary Information

This includes gel analysis of the azide-DBCO reaction between tagged protein and tagged DNA, LC-MS analysis of the azide tagging of N122C, comparisons of nanopore measurements run in duplex and triplex, comparisons of noisy and clear samples, monitoring of protein binding level for the duration of the experiment and sequences for all DNA used.

## Author Contributions

SES, RIH, UFK and MV designed research; all authors performed research, SES performed and analysed nanopore experiments, RIH performed and analysed protein azide tagging and kinetic aggregation experiments. SES, RIH and SR performed azide-DBCO coupling experiments. SES, RIH, UFK and MV wrote the manuscript.

## Conflicts of Interest

SES is funded by Oxford Nanopore. RIH is a consultant of WaveBreak Therapeutics (formerly Wren Therapeutics). SC has been a consultant of WaveBreak Therapeutics. MCC is an employee of WaveBreak Therapeutics. MV is a founder of WaveBreak Therapeutics.

