## Supplementary for "Multiplexed Digital Characterisation of Misfolded Protein Oligomers via Solid-State Nanopores"

### Methods

#### PAGE gel

Polyacrylamide gels (10% v/v, with 0.5X, pH 8, Tris-Borate-EDTA and 11 mM $MgCl_2$) were hand-cast on a PAGE loading gel setup. Details on the PAGE gels recipes can be found in **Table 1**. Once all the gel mixture additions (**Table 1**) were mixed together, 1% (w/v) APS and 0.07% (w/v) TEMED were added and the mixture was immediately vortexed and poured in between two PAGE glass slides using a Pasteur pipette. The gel comb was promptly inserted and the gel was left to polymerise for at least 45 minutes and a maximum of 1 hour. Gels were run for 120 minutes at 100 V in a running buffer containing 0.5x TBE and 11 mM $MgCl_2$. DNA was stained using GelRed® (Biotium) for 15 minutes under constant shaking. Imaging was performed using a Gel-DocIt imaging system by UPV using Visionworks software under UV excitation light and exposure times varying from 5 to 10 seconds. To check for the binding of the copper-free click chemistry reaction between DBCO-DNA and azide-labelled αS and secondarily to optimise the reaction conditions, DBCO-DNA was incubated in a 1:1 ratio (monomer : DBCO) for 1h, 3h, and overnight **(Figure S3)**.

| **Gel mixture addition** | **Amount [mL]** | **Final concentration** |
|---|---|---|
| Acrylamide/bis-acrylamide, 30 % solution | 5 | 10 % |
| 10x Tris-Borate-EDTA (TBE) | 0.75 | 0.5x |
| 0.5 M $MgCl_2$ | 0.33 | 11 mM $MgCl_2$ |
| MilliQ water | 8.92 | |
| Total volume | 15 | |
| **Polymerisation initiatiors** | | |
| 10 % (w/v) ammonium persulfate (APS) | 0.15 | 0.1 % |
| N,N,N’,N’-Tetramethylethylendiamine (TEMED) | 0.01 | 0.07 % |

**Table 1**: Recipe for a 10% PAGE gel.

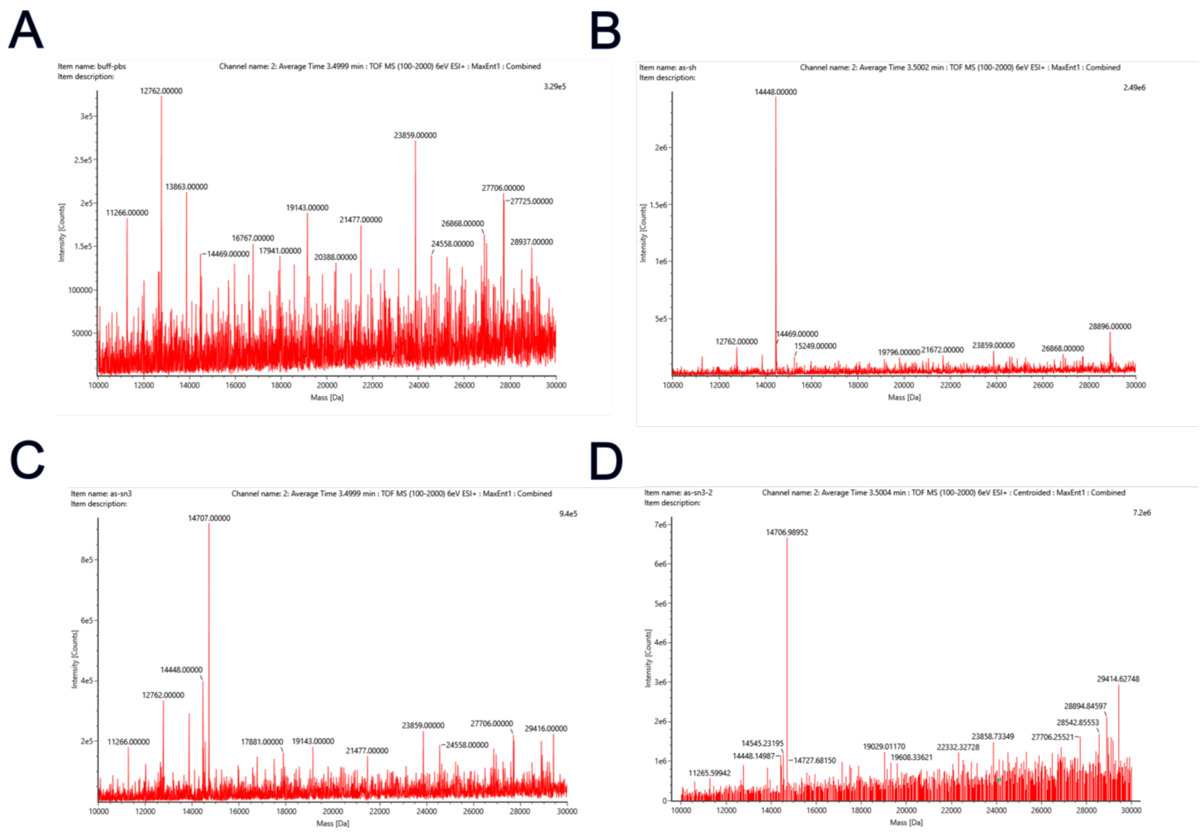

**Figure S1: LC-MS data for the reaction progress of the azide linking step to N122C-αS. (A)** PBS buffer control. (**B**) N122C (1 μM, PBS buffer) after reduction with TCEP to remove dimers, showing a single peak at 14448 Da. (**C**) Reduced N122C (1 μM, PBS) after a 2 h incubation with iodoacetamide-PEG$_3$-azide. The labelled peak, at 14707 Da, is prominent but residual unlabelled N122C remains. (**D**) After 3 h almost all of the monomer has been labelled.

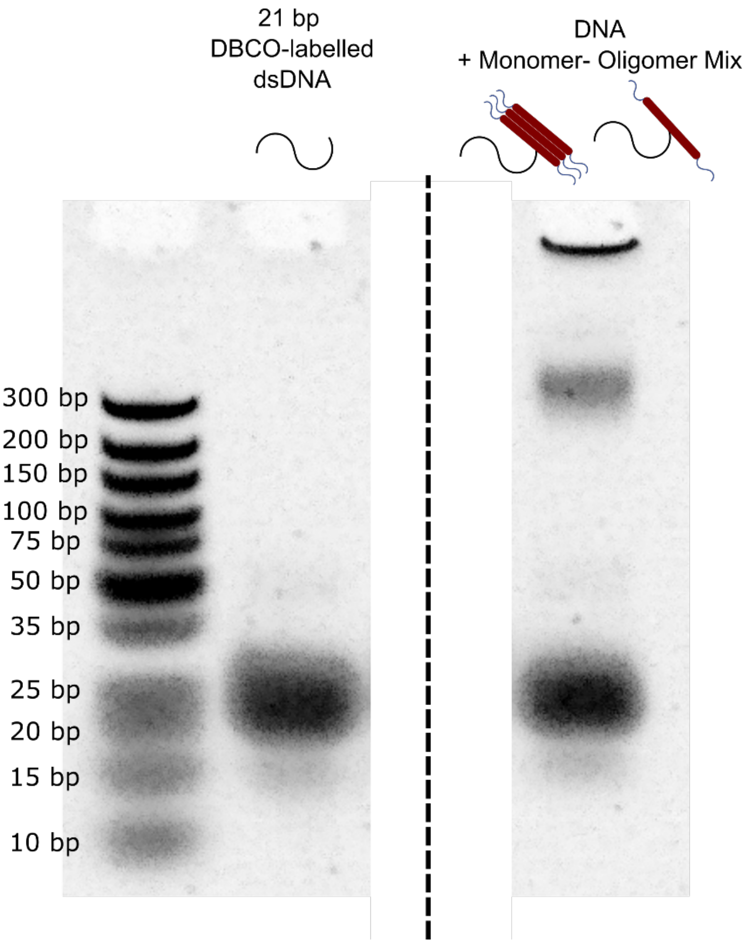

**Figure S2.** Monomer-bound DNA observed using PAGE. Column 1 shows 21 bp dsDNA and Column 2 shows a mixture of 21 bp dsDNA mixed with a partially converted oligomer and monomer sample. The monomer is 14 kDA (10 kDA ~ 270 bp), which matches the strongly stained band when added to the 21 bp DNA. The DNA retained in the well may be due to aggregates formed in the sample that cannot enter the gel.

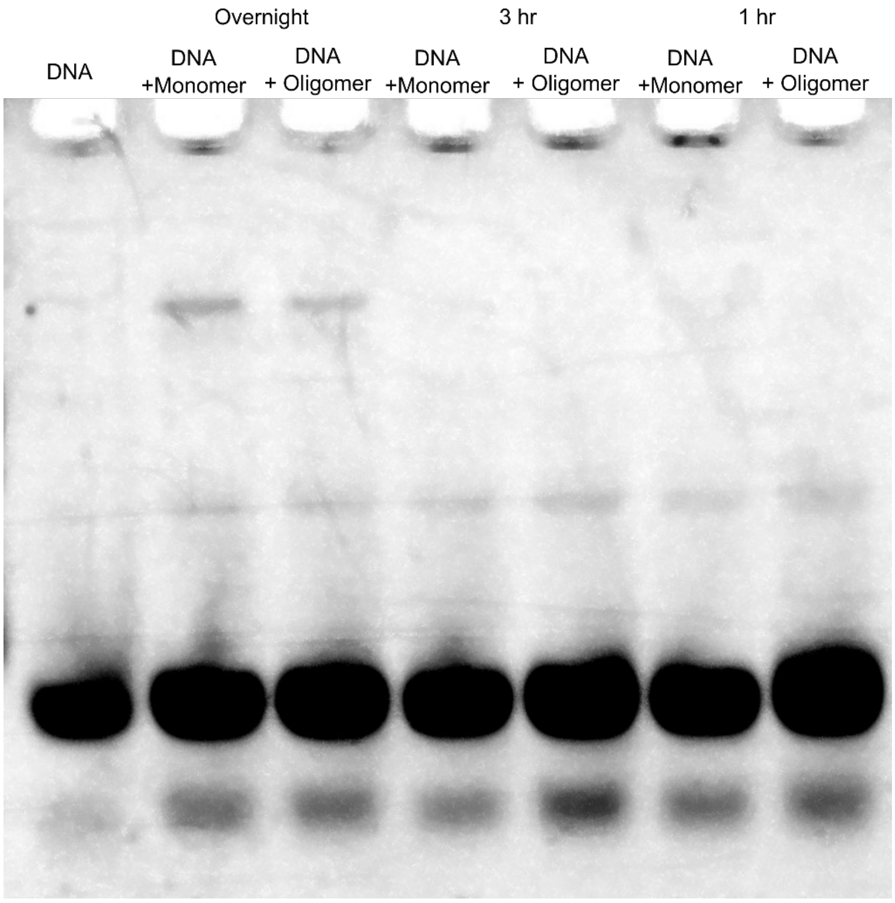

**Figure S3.** Incubation DBCO reaction observed via PAGE. Monomer is bound after overnight incubation but not observed for the 3 h and 1 h incubation times.

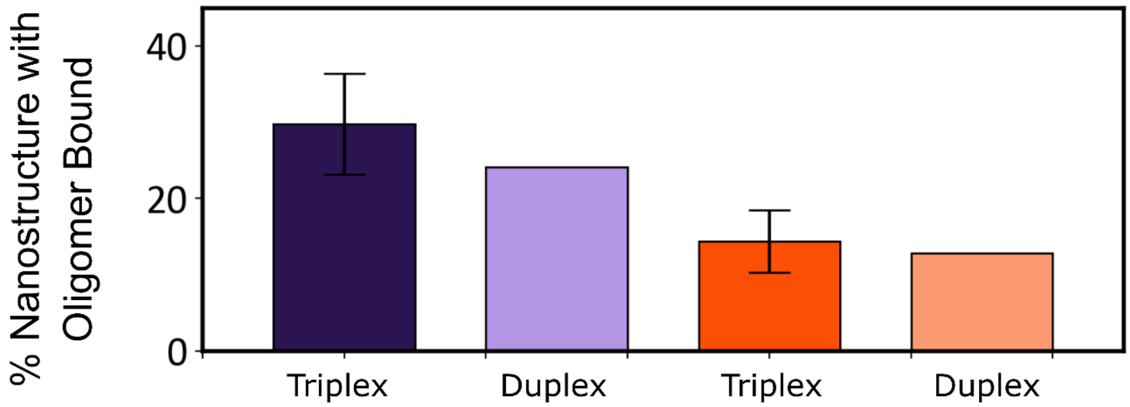

**Figure S4.** Comparison of Duplex and Triplex measurements. The samples that were measured in duplex show similar % of nanostructure with protein bound highlighting that monomer interchange is unlikely. The fraction of events with an oligomer bound to the DNA barcode; triplexed DMSO (purple) (N= 114, SD=6.62), duplexed DMSO light purple (N=54), triplexed I3.08 (orange) (N=90, SD=4.07), duplexed I3.08 (light orange) (N=39).

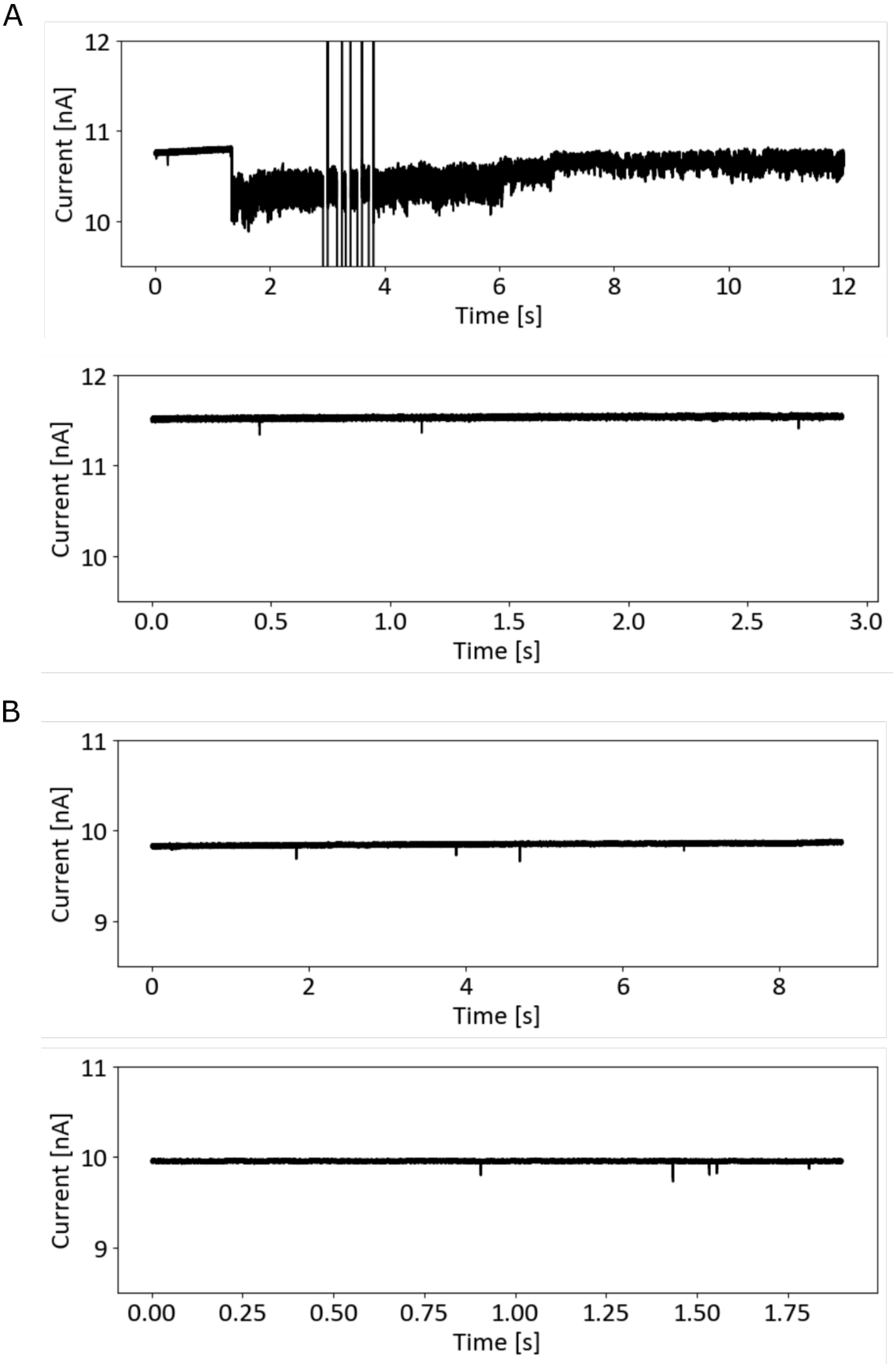

**Figure S5.** Nanopore traces with and without presence of Anle-138b. **(A)** Raw current of the nanopore trace with Anle-138b shows that upon mixture and measurement in the nanopore after 1 sec, a lot of noise is created. After 3 min (trace below) the noise level resumes back to normal. The vertical lines represent kick outs to remove protein from clogging the pore. **(B)** Raw current trace without Anle-138b shows similar noise and baseline both upon mixture and measurement in the pore and after 3 min of measurement.

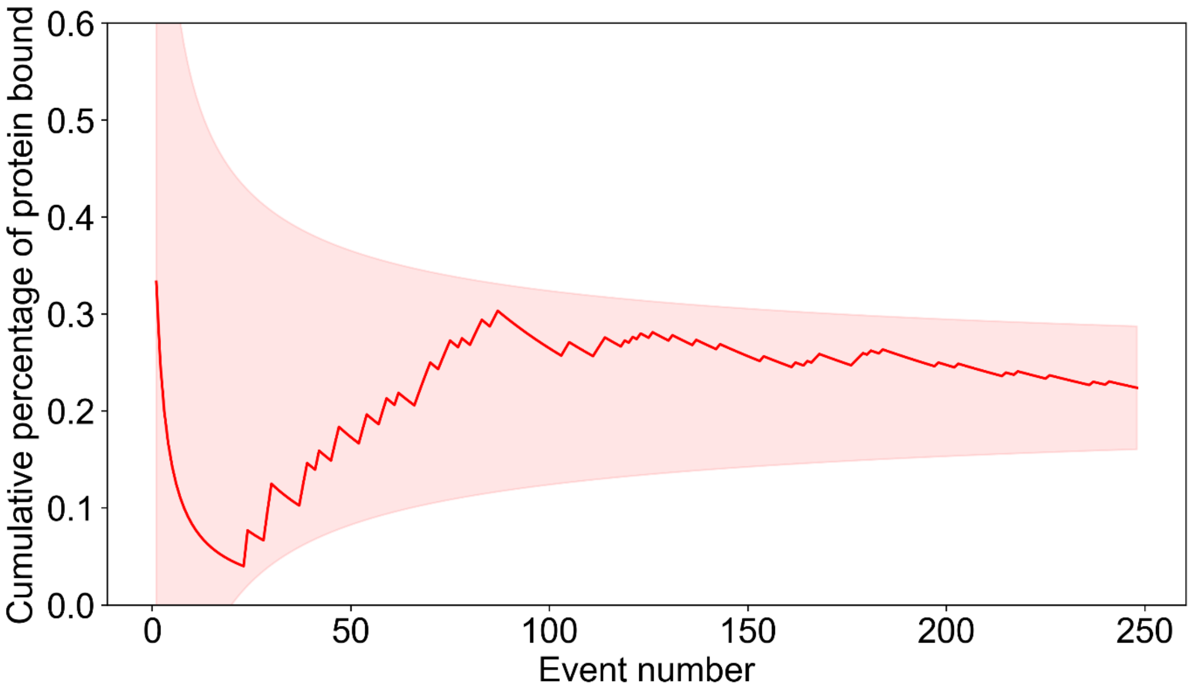

**Figure S6.** Cumulative percentage of events with clear barcode and protein bound as measured in 4 M LiCl. The number of events with the protein bound remains the same for the stabilized oligomer over the course of the 1 h measurement time. N=250 because the original trace was filtered to remove folded or knotted events with unreadable barcodes. The error fluctuation represents 1σ deviation. This becomes smaller as the number of events increases. The increase in number of events corresponds to measurement time.

The following replacements are made to create the “1” bits in the barcode portion of the nanostructure as previously shown and used in **Figure 2**[1].

First bit

Replace oligos 26,27,28,29,30,31 and 32

| |
|---|
| CTGAAAGCGTAAGAATACGTGGCACAGACAATATTTTTGAATGGCT |
| ACATCACTTGTCCTCTTTTGAGGAACAAGTTTTCTTGTCCTGAGTAGA |
| AGAACTCAAATCCTCTTTTGAGGAACAAGTTTTCTTGTCTATCGGCCT |
| TGCTGGTAATTCCTCTTTTGAGGAACAAGTTTTCTTGTATCCAGAACA |
| ATATTACCGCTCCTCTTTTGAGGAACAAGTTTTCTTGTCAGCCATTGC |
| AACAGGAAAATCCTCTTTTGAGGAACAAGTTTTCTTGTACGCTCATGG |
| AAATACCTACTCCTCTTTTGAGGAACAAGTTTTCTTGTATTTTGACGC |
| TCAATCGTCTTCCTCTTTTGAGGAACAAGTTTTCTTGTGAAATGGATT |
| ATTTACATTGTCCTCTTTTGAGGAACAAGTTTTCTTGTGCAGATTCAC |
| CAGTCACACGTCCTCTTTTGAGGAACAAGTTTTCTTGTACCAGTAATA |
| AAAGGGACATTCCTCTTTTGAGGAACAAGTTTTCTTGTTCTGGCCAAC |
| AGAGATAGAATCCTCTTTTGAGGAACAAGTTTTCTTGTCCCTTCTGAC |

Second bit

Replace oligos 40,41,42,43,44,45 and 46

| |
|---|
| AATATAATCCTGATTGTTTGGATTATACTTCTGAATAATGGAAGGG |
| CACTAACAACTCCTCTTTTGAGGAACAAGTTTTCTTGTTAATAGATTA |
| GAGCCGTCAATCCTCTTTTGAGGAACAAGTTTTCTTGTTAGATAATAC |
| ATTTGAGGATTCCTCTTTTGAGGAACAAGTTTTCTTGTTTAGAAGTAT |
| TAGACTTTACTCCTCTTTTGAGGAACAAGTTTTCTTGTAAACAATTCG |
| ACAACTCGTATCCTCTTTTGAGGAACAAGTTTTCTTGTTTAAATCCTT |
| TGCCCGAACGTCCTCTTTTGAGGAACAAGTTTTCTTGTTTATTAATTT |
| TAAAAGTTTGTCCTCTTTTGAGGAACAAGTTTTCTTGTAGTAACATTA |
| TCATTTTGCGTCCTCTTTTGAGGAACAAGTTTTCTTGTGAACAAAGAA |
| ACCACCAGAATCCTCTTTTGAGGAACAAGTTTTCTTGTGGAGCGGAAT |
| TATCATCATATCCTCTTTTGAGGAACAAGTTTTCTTGTTTCCTGATTA |
| TCAGATGATGTCCTCTTTTGAGGAACAAGTTTTCTTGTGCAATTCATC |

Third bit

Replace oligos 54,55,56,57,58,59 and 60

| |
|---|
| TGTAAATGCTGATGCAAATCCAATCGCAAGACAAAGAACGCGAGAA |

| |
|---|
| TACCTTTTTTTCCTCTTTTGAGGAACAAGTTTTCTTGTAATGGAAACA |
| GTACATAAATTCCTCTTTTGAGGAACAAGTTTTCTTGTCAATATATGT |
| GAGTGAATAATCCTCTTTTGAGGAACAAGTTTTCTTGTCCTTGCTTCT |
| GTAAATCGTCTCCTCTTTTGAGGAACAAGTTTTCTTGTGCTATTAATT |
| AATTTTCCCTTCCTCTTTTGAGGAACAAGTTTTCTTGTTAGAATCCTT |
| GAAAACATAGTCCTCTTTTGAGGAACAAGTTTTCTTGTCGATAGCTTA |
| GATTAAGACGTCCTCTTTTGAGGAACAAGTTTTCTTGTCTGAGAAGAG |
| TCAATAGTGATCCTCTTTTGAGGAACAAGTTTTCTTGTATTTATCAAA |
| ATCATAGGTCTCCTCTTTTGAGGAACAAGTTTTCTTGTTGAGAGACTA |
| CCTTTTTAACTCCTCTTTTGAGGAACAAGTTTTCTTGTCTCCGGCTTA |
| GGTTGGGTTATCCTCTTTTGAGGAACAAGTTTTCTTGTTATAACTATA |

Fourth bit

Replace oligos 68,69,70,71,72,73 and 74

| |
|---|
| TCATCGAGAACAAGCAAGCCGTTTTTATTTTCATCGTAGGAATCAT |
| AGAATATAAATCCTCTTTTGAGGAACAAGTTTTCTTGTGTACCGACAA |
| AAGGTAAAGTTCCTCTTTTGAGGAACAAGTTTTCTTGTAATTCTGTCC |
| AGACGACGACTCCTCTTTTGAGGAACAAGTTTTCTTGTAATAAACAAC |
| ATGTTCAGCTTCCTCTTTTGAGGAACAAGTTTTCTTGTAATGCAGAAC |
| GCGCCTGTTTTCCTCTTTTGAGGAACAAGTTTTCTTGTATCAACAATA |
| GATAAGTCCTTCCTCTTTTGAGGAACAAGTTTTCTTGTGAACAAGAAA |
| AATAATATCCTCCTCTTTTGAGGAACAAGTTTTCTTGTCATCCTAATT |
| TACGAGCATGTCCTCTTTTGAGGAACAAGTTTTCTTGTTAGAAACCAA |
| TCAATAATCGTCCTCTTTTGAGGAACAAGTTTTCTTGTGCTGTCTTTC |
| CTTATCATTCTCCTCTTTTGAGGAACAAGTTTTCTTGTCAAGAACGGG |
| TATTAAACCATCCTCTTTTGAGGAACAAGTTTTCTTGTAGTACCGCAC |

Fifth bit

Replace oligos 82,83,84,85,86,87 and 88

| |
|---|
| AGATAGCCGAACAAAGTTACCAGAAGGAAACCGAGGAAACGCAATA |
| AAAAATGAAATCCTCTTTTGAGGAACAAGTTTTCTTGTATAGCAGCCT |
| TTACAGAGAGTCCTCTTTTGAGGAACAAGTTTTCTTGTAATAACATAA |
| AAACAGGGAATCCTCTTTTGAGGAACAAGTTTTCTTGTGCGCATTAGA |
| CGGGAGAATTTCCTCTTTTGAGGAACAAGTTTTCTTGTAACTGAACAC |
| CCTGAACAAATCCTCTTTTGAGGAACAAGTTTTCTTGTGTCAGAGGGT |

| |
|---|
| AATTGAGCGCTCCTCTTTTGAGGAACAAGTTTTCTTGTTAATATCAGA |
| GAGATAACCCTCCTCTTTTGAGGAACAAGTTTTCTTGTACAAGAATTG |
| AGTTAAGCCCTCCTCTTTTGAGGAACAAGTTTTCTTGTAATAATAAGA |
| GCAAGAAACATCCTCTTTTGAGGAACAAGTTTTCTTGTATGAAATAGC |
| AATAGCTATCTCCTCTTTTGAGGAACAAGTTTTCTTGTTTACCGAAGC |
| CCTTTTTAAGTCCTCTTTTGAGGAACAAGTTTTCTTGTAAAAGTAAGC |

**Table S1. DNA Dumbell Bits**

| | |
|---|---|
| 142 | GATGGTTTAATTTCAACTTTAATCATTGTGAATTACCT |
| 143 | actgactgactgactgactgaTTTATGCGATTTTAAGAACTGGCTCATTATACCAGTCAGG |
| DBCO | tcagtcagtcagtcagtcagt*DBCO* |

**Table S2. DNA overhang sequences**
